## Supplemental Figures and Tables for "Microcircuit synchronization and heavy tailed synaptic weight distribution in preBötzinger Complex contribute to generation of breathing rhythm"

### Supplemental Information

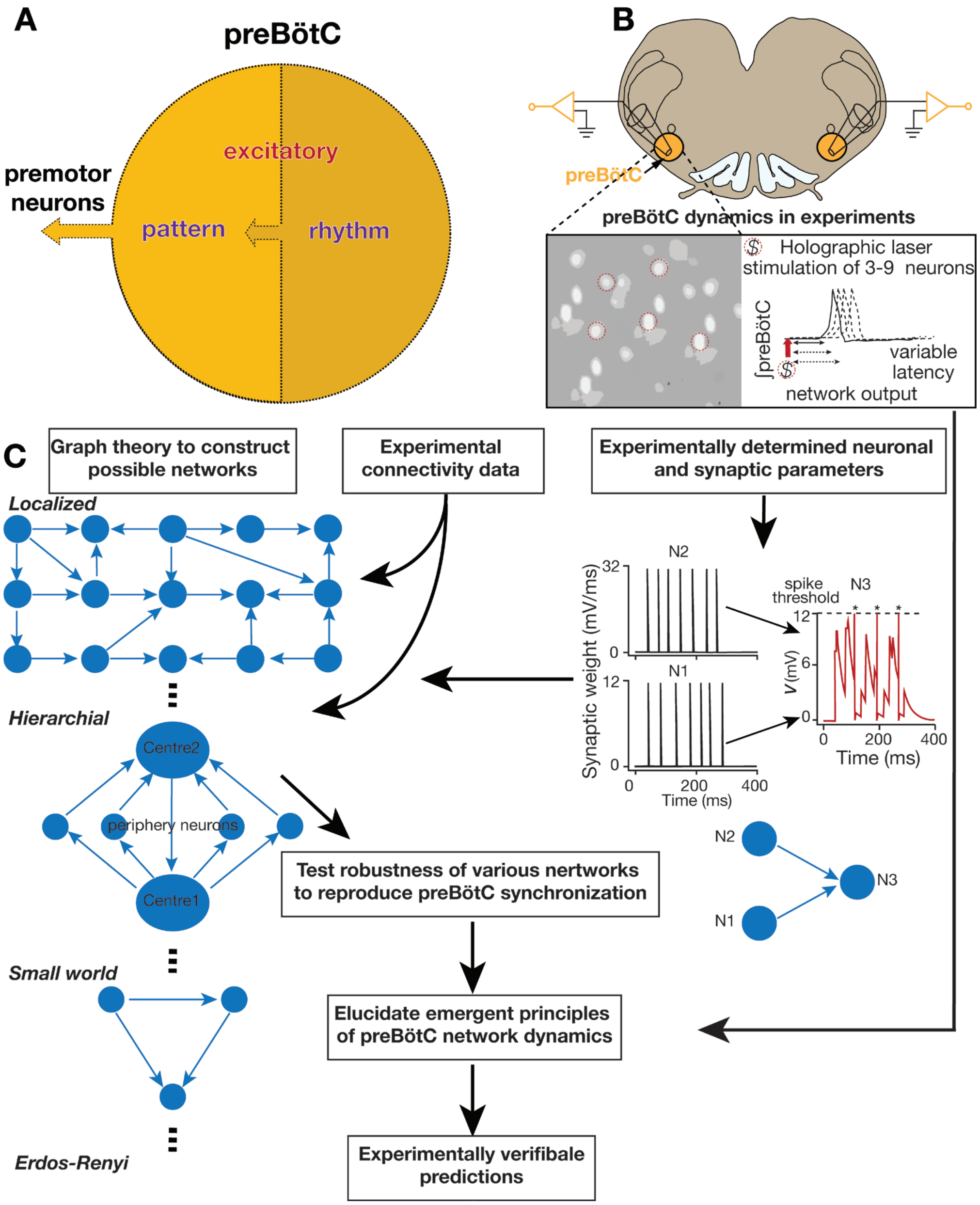


**Figure S1. Flow chart for model construction and testing.** ***Related to Figure 2*** **A**, Rhythm-generating preBötC neurons project to pattern-generating preBötC_excitatory neurons that in turn project to premotor neurons. **B**, holographic photostimulation of 3-9 preBötC inspiratory neurons (out of ~1500 total excitatory neurons) in rhythmic neonatal mouse slices generate bursts at delays ~100-500 ms (Figure 1 F-G). **C**, Various network models tested to determine if they capture preBötC dynamics depicted in **B:** (left) Network topologies as graphs with edges connected through experimentally determined connection probability between putative rhythmogenic neurons (Rekling *et.al.*, 2000)[16]. The nodes were modeled as leaky integrate and fire (LIF) neurons (right) with intrinsic and synaptic properties taken from experiments; (right) synaptic activation from two ("laser") stimulated model rhythmogenic neurons N1 and N2 (presynaptic) that project to neuron N3 (postsynaptic); their activation times represent the arrival of spikes form N1 and N2 at their respective synapses on N3 with weights indicated (left, black traces) consequently, changing in somatic potential of neuron N3 from resting potential (red, right). When the somatic potential increases above spike activation threshold $V^* = 12$ mV ($V \cong -48$ mV), N3 generates an action potential (*), followed by its potential dropping to $V_{rest} = 0$ mV ($V_{rest} \cong -60$ mV) for refractory period of 3 ms.

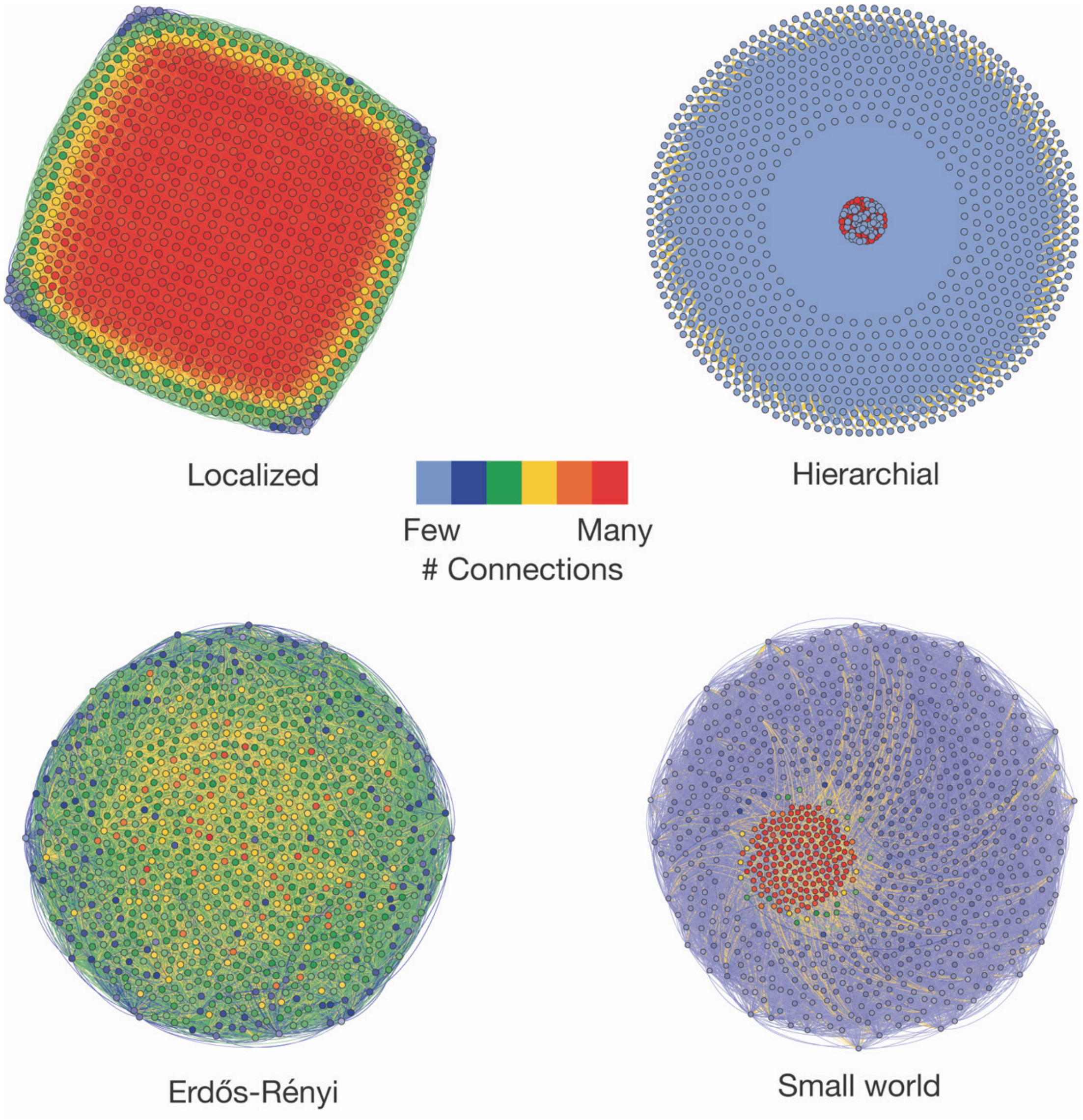


**Figure S2.** ***Related to Figure 2.*** Various networks represented on a force-based (Fruchterman-Reingold algorithm in the *Gephi* software) layout. linked nodes (neurons) are *pulled* together and unrelated nodes are farther apart; most strongly linked nodes (through direct connections or common inputs) are at the center and the least linked ones at the periphery. Nodes (and their edges) are color coded based on number of their projections, i.e., outward synapses with warmer shades representing more connections. These figures are enlarged versions of the network layouts presented in Figure 2.

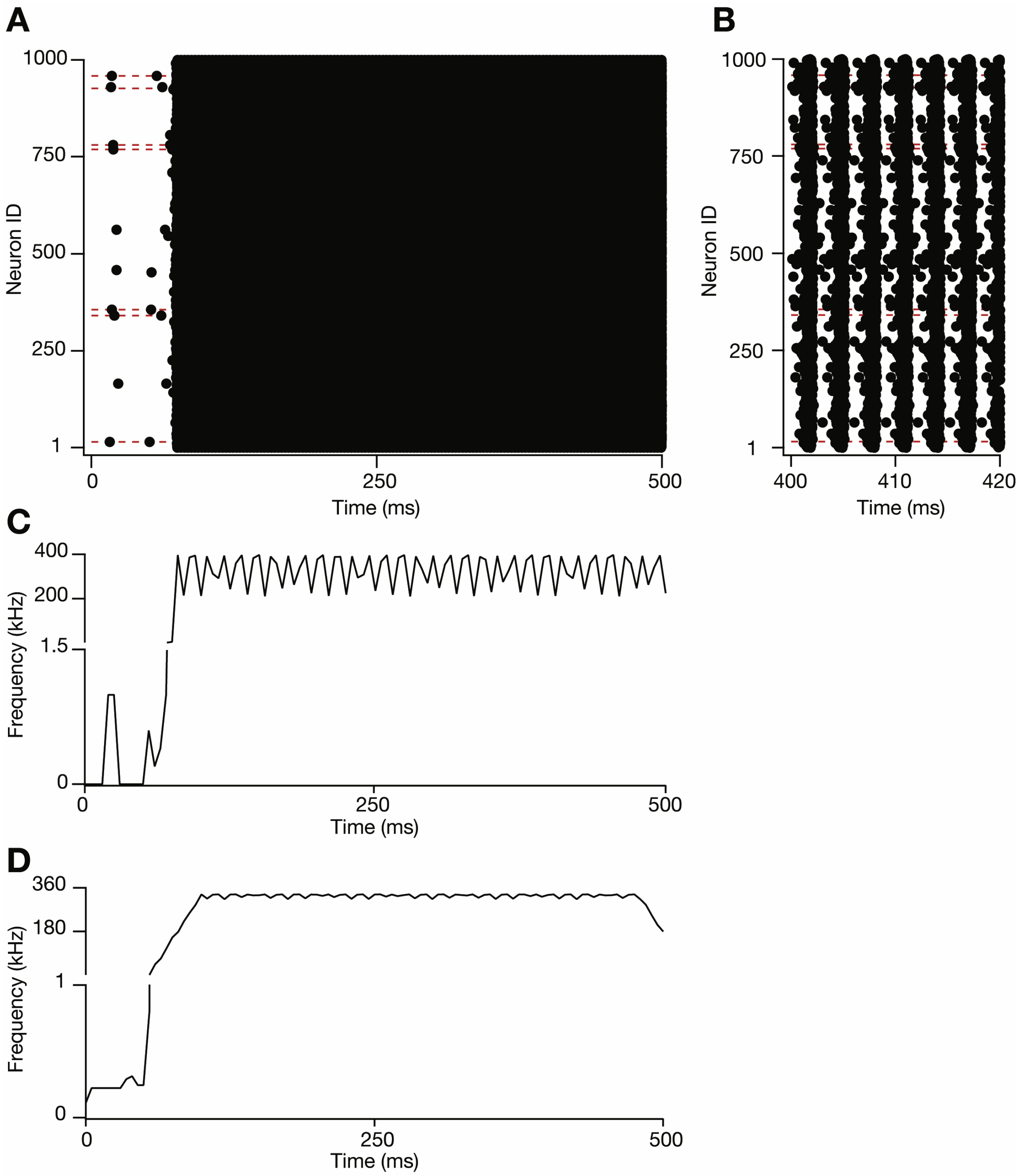


**Figure S3**. ***Related to Figure 4.*** **A**, Model output when the same set of randomly selected 7 neurons (on the dashed lines) was stimulated to fire seven spikes each at 25 Hz with 5 ms jitter, i.e., SD to model holographic uncaging of glutamate onto these neurons as in (Kam et. al., 2013)[30.] Each dot represents the time of spike (abscissa) from corresponding neuron (ordinate), note that once the network synchronized it continues in the high-frequency firing, i.e., bursting mode even after the stimulated spikes, from the seven neurons, ended at ~300 ms. **B**, Enlarged region from (a) exhibiting firing rate modulation due to the refractory period, of 3 ms, in the model neurons. **C**, average firing rate of network computed by averaging network activity in a moving window of 40 ms with 5 ms step increment.

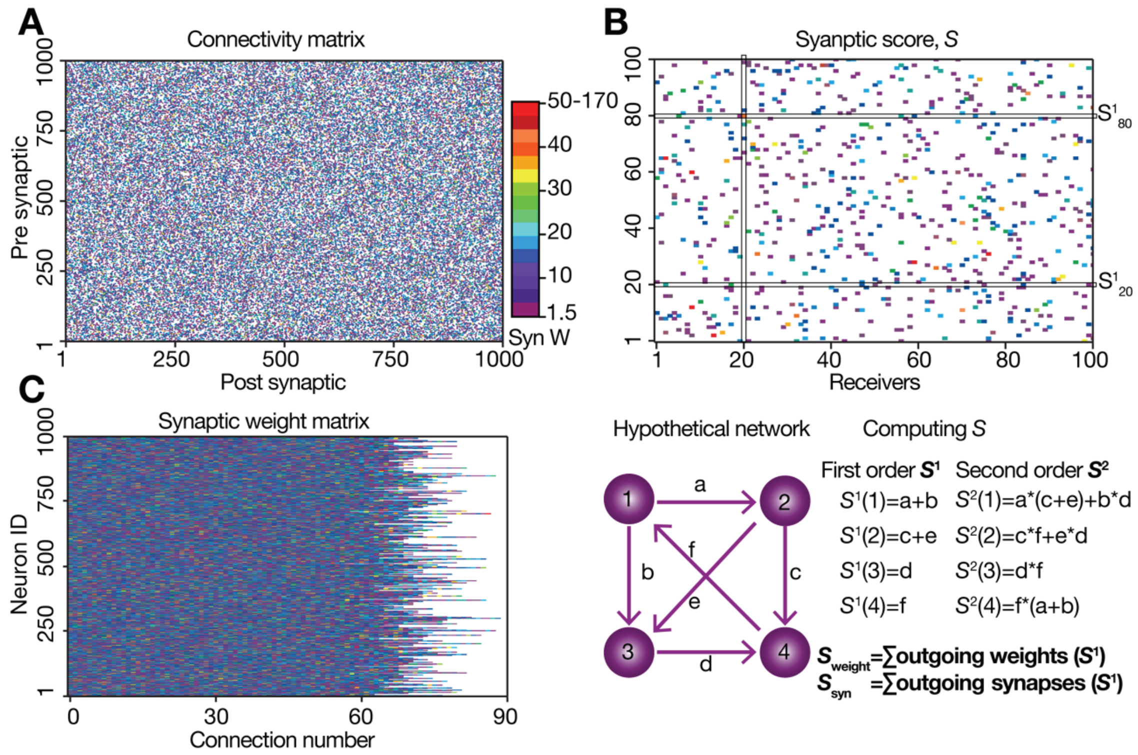


**Figure S4. Computing efferent synaptic score from network topology**. ***Related to Figures 5-6***.The efferent synaptic score, *S*, is neuron property defined by its network connectivity. **A,** Connectivity matrix of ER network with lognormal (LN) weight distribution. Presynaptic neurons (ordinate) project to other neurons labelled as postsynaptic (abscissa) with synaptic weights, randomly drawn from a LN distribution. Each dot represents synaptic weight (Syn W; color-coded) of presynaptic #i (1-1000) to postsynaptic #j (1-1000). B, top, expanded section from A (100x100) showing contribution of synaptic connections towards *S*. For example, the first order *S* of neuron #80 ($S^1_{80}$) is the sum of all synaptic weights with which it connects to other neurons. In this example, neuron #20 is postsynaptic to neuron #80, thus, $S^1_{20}$ contributes to the second order *S* of neuron #80 ($S^2_{80}$) (Second order *S* ($S^2$) also accounts for the connectivity of the next neighbors of a given neuron, see text for the calculation of $S^2$); bottom, example of *S* calculation in a hypothetical network. For a given neuron, $S_{weight}$ is the sum of its efferent synaptic weights and $S_{synout}$ is the number of its efferent synaptic connections. **C**, plot of number of outward connections of neurons in ER network with their synaptic weights color-coded as in A and B. This network was used in Figure 5; *S* varies from neuron to neuron as their outward synaptic connections vary, per the ER graph, and as their output synaptic weights vary due to the LN weight distribution.

### Supplemental Tables

### Table S1. Model parameters

| Parameter | Value |
|---|---|
| $V_{rest}$ | -60 mV; -65 mV (figure7) |
| V* | -48 mV |
| $\tau_m$ | 25 ms |
| $\tau_s$ | 0.5 ms |
| $\Delta t_{ij}$ | 1.3±1.1 ms |
| $W_{ij}$ | 300±160 (mean ± SD) mV/ms =15±8 in step size (0.05 ms) |
| $\tau_{delay}$ | 20±3 ms |
| $T_{laser}$ | 39±5 ms |
| $n_{spikes}$ | 7 |
| $f_{noise}$ | 0.5-2 Hz |

**Table S2. Prediction accuracy of various generalized *S* quantities for network burst**

| Parameter | Accuracy | Standard deviation |
|---|---|---|
| $\Sigma\ (W*Y^3)^2$ | 0.73 | 0.01 |
| $\Sigma\ W*Y^4$ | 0.72 | 0.02 |
| $\Sigma\ (W*Y^3)^3$ | 0.72 | 0.006 |
| $\Sigma\ (W*Y^4)^2$ | 0.72 | 0.009 |
| $\Sigma\ Y^4$ | 0.72 | 0.01 |
| $\Sigma\ (W*Y^4)^3$ | 0.71 | 0.01 |
| $\Sigma\ Y^4$ | 0.71 | 0.02 |
| $\Sigma\ Y^6$ | 0.71 | 0.007 |
| $\Sigma\ Y^6$ | 0.70 | 0.006 |
| $\Sigma\ W*Y^3$ | 0.70 | 0.01 |
| $\Sigma\ Y^3$ | 0.70 | 0.01 |
| $\Sigma\ Y^3$ | 0.70 | 0.01 |
| $\Sigma\ (W*Y^2)^3$ | 0.70 | 0.01 |
| $\Sigma\ Y^9$ | 0.69 | 0.01 |
| $\Sigma\ Y^8$ | 0.69 | 0.008 |
| $\Sigma\ Y^{12}$ | 0.68 | 0.01 |
| $\Sigma\ (W*Y^2)^2$ | 0.68 | 0.009 |
| $\Sigma\ W*Y^2$ | 0.66 | 0.009 |
| $\Sigma\ Y^2$ | 0.66 | 0.02 |
| $\Sigma\ Y^2$ | 0.66 | 0.01 |
| $\Sigma\ (W*Y)^3$ | 0.59 | 0.008 |
| $\Sigma\ Y$ | 0.57 | 0.02 |
| $\Sigma\ (W*Y)^2$ | 0.57 | 0.01 |
| $\Sigma\ W*Y$ | 0.56 | 0.01 |
| *All above simultaneously* | 0.74 | 0.006 |
